# Lift the Veil of Breast Cancers Using Four or Fewer Critical Genes

**DOI:** 10.1101/2021.10.19.465026

**Authors:** Zhengjun Zhang

## Abstract

Tackling breast cancer problems is like mastering a puzzle, and the mystery is not yet solved. Reported key genes in the literature could not be confirmed whether they are vital to breast cancer formations due to lack of convincing accuracy, although they may be biologically directly related to breast cancer based on present biological knowledge. It is hoped vital genes can be identified with the highest possible accuracy, e.g., 100% accuracy and convincing causal patterns beyond what has been known in breast cancer. One hope is that finding gene-gene interaction signatures and functional effects may solve the puzzle. This research uses a recently developed competing linear factor analysis method in differentially expressed gene detection to advance the study of breast cancer formation to its deepest root level as deep as possible. Surprisingly, three genes are detected to be differentially expressed in TNBC, and non-TNBC (Her2, Luminal A, Luminal B) samples with 100% sensitivity and 100% specificity in one study of triple-negative breast cancers (TNBC, with 54675 genes and 265 samples). These three genes show a clear signature pattern of how TNBC patients can be grouped. For another TNBC study (with 54673 genes and 66 samples), four genes bring the same accuracy of 100% sensitivity and 100% specificity. Four genes are found to have the same accuracy of 100% sensitivity and 100% specificity in one breast cancer study (with 54675 genes and 121 samples), and the same four genes bring an accuracy of 100% sensitivity and 96.5% specificity in the fourth breast cancer study (with 60483 genes and 1217 samples.) These results show the four-gene-based classifiers are robust and accurate. The detected genes naturally classify patients into subtypes, e.g., seven subtypes. These findings demonstrate the clearest gene-gene interaction patterns and functional effects with the smallest numbers of genes and the highest accuracy compared with findings reported in the literature. The four genes are considered to be essential for breast cancer studies and practice. They can provide focused, targeted researches and precision medicine for each subtype of breast cancer. New breast cancer disease types may be detected using the classified subtypes, and hence new effective therapies can be developed.

## Introduction

Breast cancer has been an unconquered plague for centuries. It has had the highest death rate among all cancers women have had for many years. It has caused enormous economic losses and costs. To save lives and protect women from breast cancers, enormous research efforts and money have been investigated. Although there have been some considerable signs of progress in breast cancer diagnoses and therapies, many women still suffer from being diagnosed with breast cancer and lost their lives every year. No apparent clues or research results show the most critical genetic causality in breast cancer formation. The most hopeful direction, finding critical genes, or primary differentially expressed genes related to breast cancer formation, has been drawing much attention in breast cancer studies. The most recent editorial summary by Narod (2021) states “Results of two large case-control studies that analyzed the associations between a number of putative cancer susceptibility genes and breast cancer risk are now reported in the Journal. The study by Dorling et al. (2021, Breast Cancer Association Consortium) included 34 genes and 113,000 women from 25 countries, and the study by Hu et al. (2021) included 28 genes and 64,000 women from the United States. Variants in 8 genes — BRCA1, BRCA2, PALB2, BARD1, RAD51C, RAD51D, ATM, and CHEK2 — had a significant association with breast cancer risk in both studies.”

Differential expression analysis between tumor and non-tumor cells helps breast cancer prognosis prediction at a relatively early stage, identifying some clear patterns from patients to patients, recommending different precision therapies according to breast cancer subtypes. Efforts have been made in identifying genes associated with breast cancer symptoms. For example, in a systems biology comprehensive analysis on breast cancer to identify key gene modules and genes associated with TNM-based clinical stages (Amjad et al. 2020), the authors have identified various numbers of genes that can be key genes related to breast cancers at different cancer stages. Malvia et al. (2019) studied gene expression profiles of breast cancers in Indian women, obtained 2413 differentially expressed genes, and demonstrated the existence of molecular subtypes in Indian women. Lv et al. (2019) aimed to explore some novel genes and pathways related to TNBC prognosis through bioinformatics methods as well as potential initiation and progression mechanisms. 755 differentially expressed overlapping mRNAs were detected between TNBC/non-TNBC samples and normal tissue. The authors found eight hub genes associated with the cell cycle pathway highly expressed in TNBC. Additionally, a novel six-gene (TMEM252, PRB2, SMCO1, IVL, SMR3B, and COL9A3) signature from the 755 differentially expressed mRNAs were constructed and significantly associated with prognosis as an independent prognostic signature. Zhong et al. (2020) conducted a robust rank aggregation (RRA) analysis based on genome-wide gene expression datasets involving TNBC patients from the Gene Expression Omnibus (GEO) database to identify key genes associated with TNBC. A total of 194 highly ranked differentially expressed genes (DEGs) were identified in TNBC vs. non-TNBC. Gene oncology (GO) and Kyoto Encyclopedia of Genes and Genomes pathway (KEGG) enrichment analysis was utilized to explore the identified genes’ biological functions. The authors also found that some genes are positively correlated to the life expectancy (P<0.05) of TNBC patients. Lin et al. (2020) identified potential key genes for HER-2 positive breast cancer based on bioinformatics analysis. A total of 54 up-regulated DEGs and 269 downregulated DEGs were identified. Among them, 10 hub genes including CCNB1, RAC1, TOP2A, KIF20A, RRM2, ASPM, NUSAP1, BIRC5, BUB1B, and CEP55 demonstrated by connectivity degree in the PPI network were screened out. Chen et al. (2018) systematically searched the electronic databases of MEDLINE (PubMed), Embase, and Cochrane Library to identify relevant publications from April, 1959 to November, 2017. identified 16 qualified studies from 527 publications with 46,870 breast cancer patients including 868 BRCA1 mutations carriers, 739 BRCA2 mutations carriers, and 45,263 non-carriers. The results showed that breast cancer patients with BRCA1Mut carriers were more likely to have TNBC than those of BRCA2Mut carriers (OR: 3.292; 95% CI: 2.773–3.909) or non-carriers (OR: 8.889; 95% CI: 6.925–11.410). Deng et al. (2019) identified potential crucial genes and key pathways in breast cancer using bioinformatic analysis. 203 up-regulated and 118 down-regulated DEGs were identified. Six hub genes were selected and validated in clinical sample for further analysis due to the high degree of connectivity, including CDK1, CCNA2, TOP2A, CCNB1, KIF11, and MELK. They were all correlated to worse overall survival (OS) in breast cancer. Zhu et al. (2020) identified some key genes and pathways associated with irradiation in breast cancer tissue and breast cancer cell lines. A total of 82 DEGs (74 up-regulated and 8 downregulated genes) were identified. Two characteristic subnetworks and 3 hub genes (FOS, CCL2, and CXCL12) were strongly distinguished in PPI network. Dong et al. (2018) aimed to identify the key pathways and genes and find the potential initiation and progression mechanism of TNBC. 56 up-regulated and 151 downregulated genes were listed and the gene oncology (GO) and Kyoto Encyclopedia of Genes and Genomes pathway (KEGG) enrichment analysis was performed. The authors found that SOX8, AR, C9orf152, NRK and RAB30, and other key genes and pathways might be promising targets for the TNBC treatment. Lu et al. (2019) identified five hub genes (PHLPP1, UBC, ACACB, TGFB1, and ACTB) associated with HER2+BC with brain metastasis. The GSEA analysis revealed that the ribosomal pathway seems to play a very important role in the pathogenesis of HER2+BC with brain metastasis. Such studies are many. This paper won’t be able to review all of them. Readers can search and find interesting topics online.

The reported genes in the published work point out some promising directions in breast cancer research and treatments. But it is not clear whether or not they are fundamental causes or direct causes of breast cancer. The problem is mainly due to the following three main limitations. 1) The number of human genes is ultra-large compared to the number of patients in affordable study designs. Identifying a few key (single digit) genes that are uniformly optimal across different trials, different study purposes, different measurement methods, and different cohorts is rather challenging. From the aforementioned research outcomes, we can see there are many different genes are identified. As a result, it’s impossible to see which one is the most important one, which can be a driver of breast cancer disease. 2) The inefficient detecting power of existing analysis methods due to restricted model assumptions cannot deal with heterogeneous populations (different breast cancer subtypes). As a result, the sensitivity and specificity of many published gene classifiers are not satisfactory. 3) It isn’t easy to extract informative messages from existing models and analysis methods. Also, many gene-related classifiers are not interpretable as gene-gene inter-relationships and functional effects are hardly expressed. As a result, scientific research progress in breast cancer studies is still limited. Much literature attention has been focused on individual genes and their expression levels, i.e., not gene-gene interactions, genes-subtypes (of breast cancers) interactions, and functional effects. As a result, the fundamental genetic causes of breast cancer formations can be masked by those suboptimal focuses, and the researches can still be in a primitive state. Many unknown factors exist. They can be essential to conquer the breast cancer plague, and therefore there is an urgent need for identifying critical DEGs with the highest possible sensitivity and specificity for breast cancer detection.

This work aims to lift the veil of breast cancers by discovering the joint functional effects of four or fewer critical DEGs that show the highest detecting power of breast cancer in four gene expression RNA-seq datasets. According to our analysis, these four genes and their functional effects describe breast cancers’ overall features at the genomic level, with the highest possible sensitivity (up to 100%) and specificity (up to 100%) for breast cancer detection. In addition, they are invariance preserving with the same group of patients but measured in different scales, and they are robust from one trial to another trial.

## Statistical Methodology

The most recently developed max-linear competing factor models (Cui and Zhang, 2018), max-linear regression models (Cui et al. 2020), and max-linear logistic models (Xu 2019, Zhang 2021) have proven to be powerful models and analysis approaches to study heteroscedastic populations and competing risks and resources. The theoretical foundations of these models have been established in Cui and Zhang (2018), Cui et al. (2020), Malinowski et al. (2018), Xu (2019), and Zhang (2020, 2021). The difference between the max-linear competing models and the classical statistical models is that the original linear combination of predictors is replaced by the maximum of a set of linear combinations of predictors, called competing factors or competing-risk factors. The max-linear competing factor models are different from existing popular classification models such as random forest, support vector machine, group lasso-based machine learning methods, and deep learning methods. However, the max-linear competing factor models are interpretable and outperform existing methods (Cui et al. 2020). This study implements the max-linear logistic regression model to build a competing factor breast cancer classifier. For completeness, the model is stated as follows.

Suppose there are *i* = 1, …, *n* patients with breast cancer status label *Y*_*i*_ = 1 for cancer and *Y*_*i*_ = 0 for cancer-free, and *Y*_*i*_ is related to *G* groups of genes by

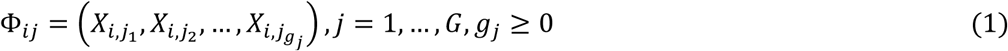

where *i* is the *i*th individual in the sample, *g*_*j*_ is the number of genes in *j*th group. The competing (risk) factor classifier for the *i* th outcome variable is defined as

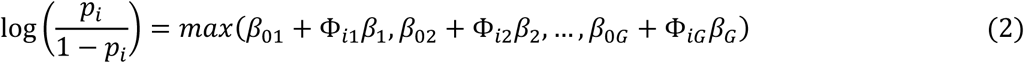

where *β*_0*j*_’s are intercepts, Φ_*ij*_ is a 1 × *g*_j_ observed vector, *β*_j_ is a *g*_j_ × 1 coefficient vector which characterizes the contribution of each predictor to the outcome variable **Y** in the *j*th group to the risk, and *β*_0*j*_ + Φ_*ij*_*β*_*j*_ is called the *j*th competing risk factor, i.e., *j*th signature. The unknown parameters are estimated from

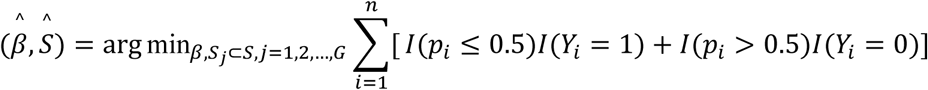

where 0.5 is a probability threshold value that is commonly used in machine learning classifiers, *I*(.) is an indicate function, *p*_*i*_ is defined in the equation (2). *S* = {1,2, …,54675}is the index set of all genes,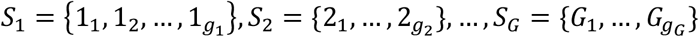 are index sets corresponding to (1), and 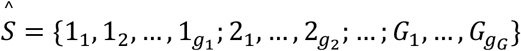 is the final gene set selected in the final classifiers.

The goal is to identify the clearest patterns of gene-gene interactions and functional effects related to breast cancer samples and non-tumor samples. We start with three competing factors in max-linear logistic regression models, with each factor having only three genes randomly drawing from 54675, 54673, or 60483 genes. Then, a Monte Carlo method with extensive computation is used to find the final model with the best performance of sensitivity and specificity and the smallest number of genes. Finally, the complete computing description is listed in Zhang (2021) in which five Covid-19 critical genes and seven subtypes were identified.

## Data Descriptions

There are four datasets used in this study. The first dataset is triple-negative breast cancer (TNBC, North American cohort) study conducted by Burstein et al. (2015) and den Hollander (2016) with 54675 genes, 198 TNBC tumor samples, and 67 not TNBC (Her2, Luminal A, Luminal B) samples. The data link and descriptions are https://www.ncbi.nlm.nih.gov/geo/query/acc.cgi?acc=GSE76275. The platforms are GPL570 [HG-U133_Plus_2] Affymetrix Human Genome U133 Plus 2.0 Array. The expression values are log2(RMA signal). The second dataset is a European cohort with 55 TNBC samples and 11 normal breast tissue samples, Maire et al. (2013), Maire et al. (2013), Maubant et al. (2015). The data link and description are https://www.ncbi.nlm.nih.gov/geo/query/acc.cgi?acc=GSE65194. The platforms are GPL570 [HG-U133_Plus_2] Affymetrix Human Genome U133 Plus 2.0 Array. The number of genes is 54673. The expression values are log2(GCRMA signal from Affy cdf). The third dataset is gene expression profiling of 104 breast cancer and 17 normal breast biopsies by Clark et al. (2013). It is from a European cohort. The data link and descriptions are https://www.ncbi.nlm.nih.gov/geo/query/acc.cgi?acc=GSE42568. The platforms are GPL570 [HG-U133_Plus_2] Affymetrix Human Genome U133 Plus 2.0 Array. The expression values are log2(GC-RMA signal intensity). The fourth dataset is GDC TCGA Breast Cancer cohort by Genomic Data Commons. The dataset contains 60,484 identifiers (genes) and 1217 (1104 tumors and 113 tumor free) samples. Data from the same sample but from different vials/portions/analytes/aliquotes is averaged; data from different samples are combined into genomicMatrix; all data is then log2(fpkm+1) transformed. The platform is Illumina. The type of data is gene expression RNAseq. The data link and descriptions are https://xenabrowser.net/datapages/?dataset=TCGA-BRCA.htseq_fpkm.tsv&host=https%3A%2F%2Fgdc.xenahubs.net&removeHub=https%3A%2F%2Fxena.treehouse.gi.ucsc.edu%3A443.

## Results and Interpretations

Using a probability higher than 50% as the threshold, we identify three critical DEGs: RBM22 (RNA binding motif protein 22), RNF213 (ring finger protein 213), and CACNG4 (Calcium Voltage-Gated Channel Auxiliary Subunit Gamma 4), which lead to 100% sensitivity and 100% specificity of classifying all 265 samples in their respective groups in the first TNBC dataset; four critical DEGs: MYCT1 (MYC Target 1), NUAK2 (NUAK Family Kinase 2), NAT8L (N-Acetyltransferase 8 Like), and CACNG4, which lead to 100% sensitivity and 100% specificity of classifying all 66 samples in their respective groups in the second TNBC dataset; four critical DEGs: MYCT1, UNC5B (Unc-5 Netrin Receptor B), NUAK2, and NAT8L, which also lead to 100% sensitivity and 100% specificity of classifying all 121 samples in their respective groups in the third breast cancer dataset; and the same four critical DEGs as in the third dataset, which leads to 100% sensitivity and 96.5% specificity of classifying all 1217 samples in their respective groups in the fourth breast cancer dataset.

Our final classifiers are combined classifiers of three competing factor (CF*i, i*=1,2,3) classifiers expressed as:

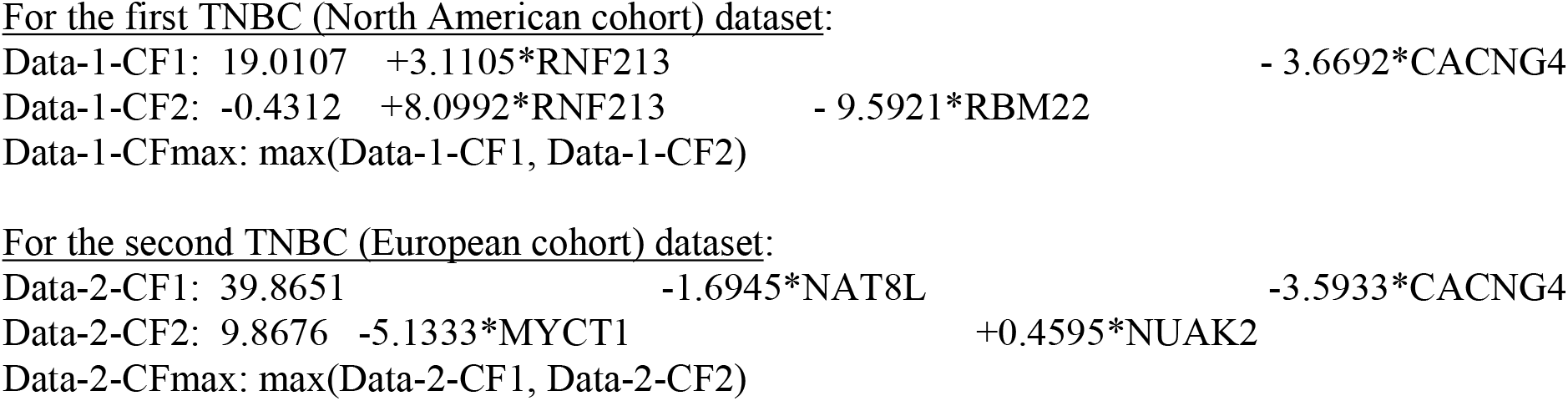
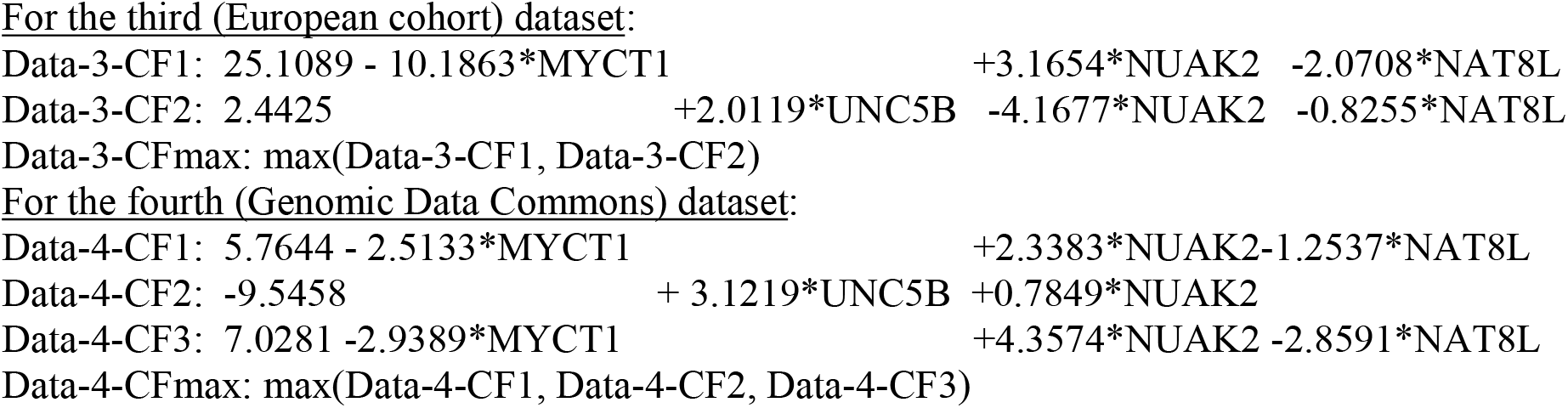

The risk probabilities are calculated using the logistic function of exp(Data-*i*-CFmax)/(1+ exp(Data-*i*-CFmax)) for the combined classifiers in each dataset, or exp(Data-*i*-CF*j*)/(1+ exp(Data-*i*-CF*j*)) for each individual classifier *i*=1,2,3, 4, *j*=1,2,3.

In the first three cohorts, multiple ID-ref subtype genes correspond to a gene symbol. The following ID-ref subtype genes are used in the classifiers: 236872_at (RBM22), 241480_at (RNF213), 62987_r_at (CACNG4), 220471_s_at (MYCT1), 220987_s_at (NUAK2), 228880_at (NAT8L), 226899_at (UNC5B).

Table 1 lists gene expression values, individual classifiers’ computed values, the combined classifier’s computed values, and the risk probabilities. Figure 1 plots all patients’ risk probabilities with circles for breast cancer samples and asters for non-breast cancer samples. Figure 2 is a Venn diagram that plots individual classifiers’ performance. This study is the first time TNBC and other breast cancer types can be further classified into subtypes based on critical genes’ functions. This new classification opens a new research direction, new drug developments, and new refined personalized therapies.

**Table 1.**
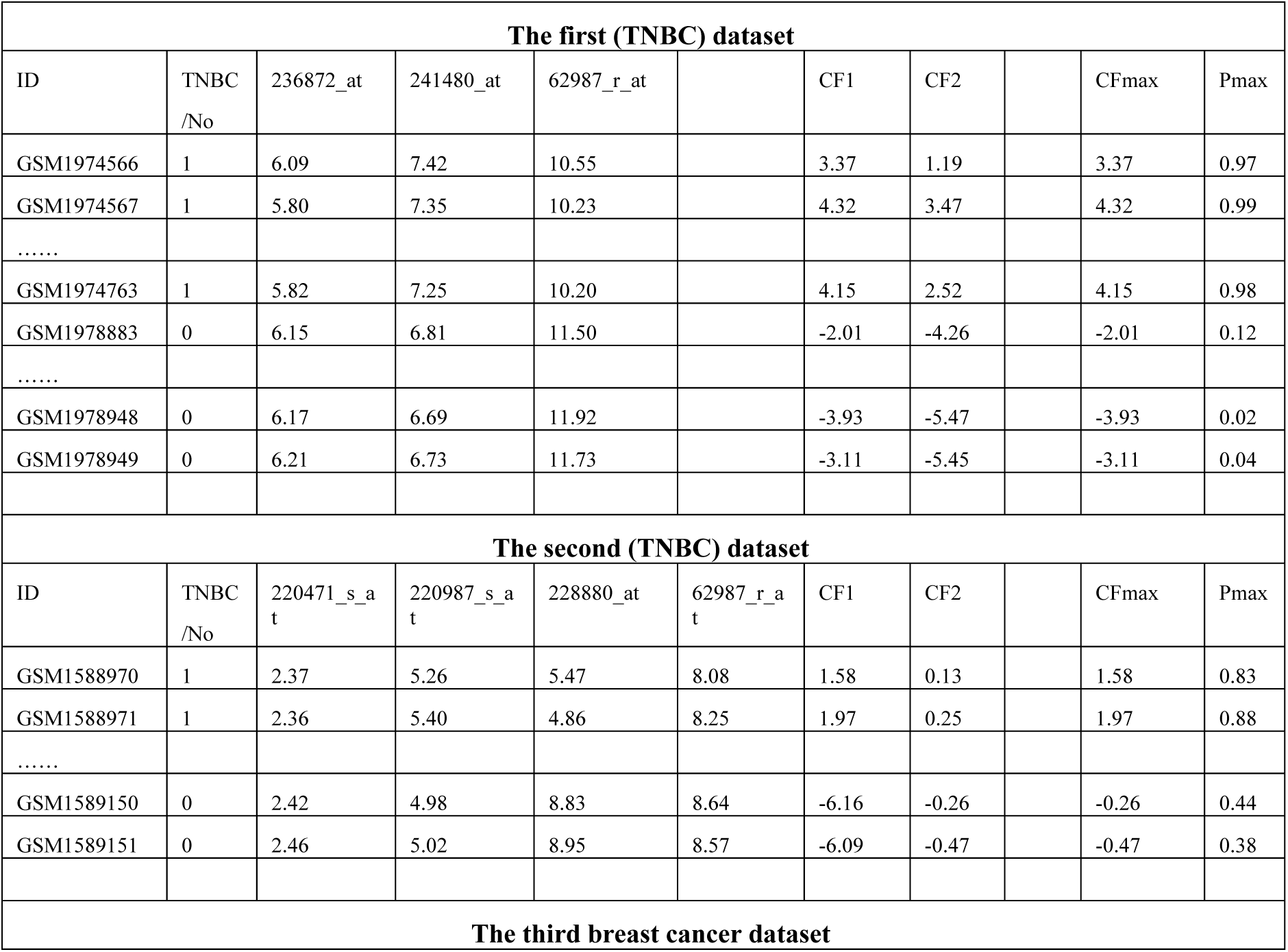
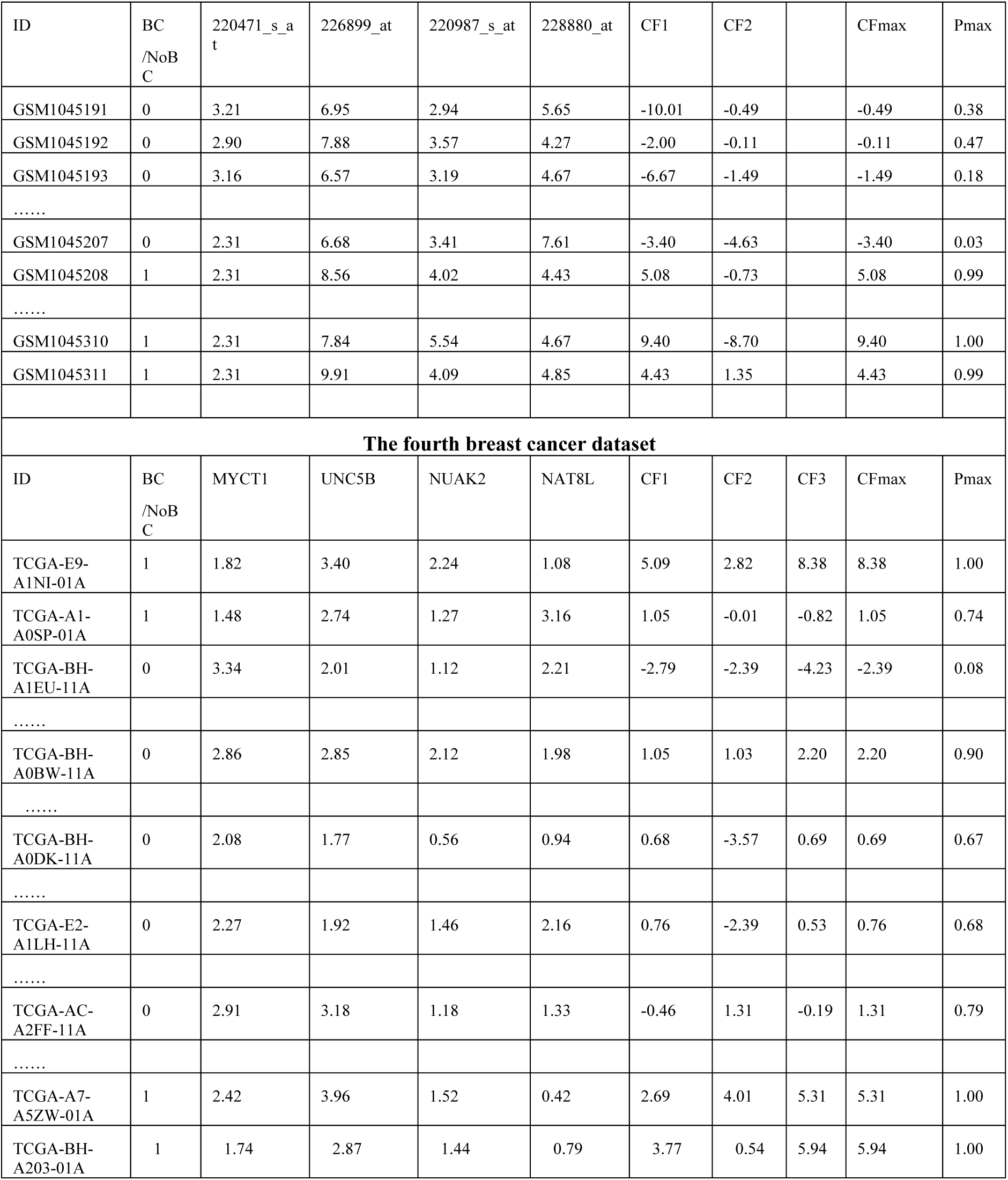
Gene information, expression values, competing factors, risk probabilities.

**Figure 1.**
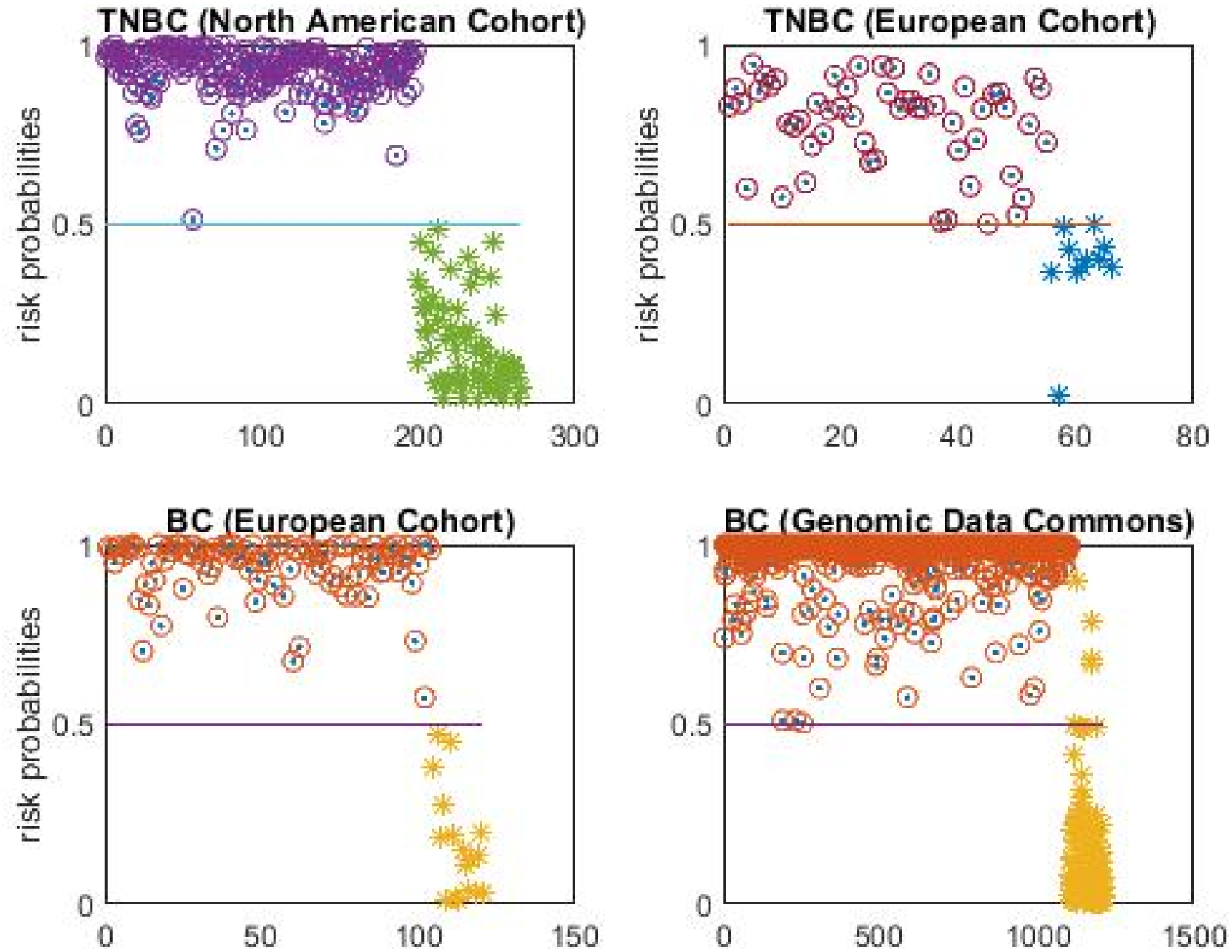
Risk probabilities of four cohorts. The circles are for patients with breast cancers. The asters are for tissues without breast cancers.

**Figure 2.**
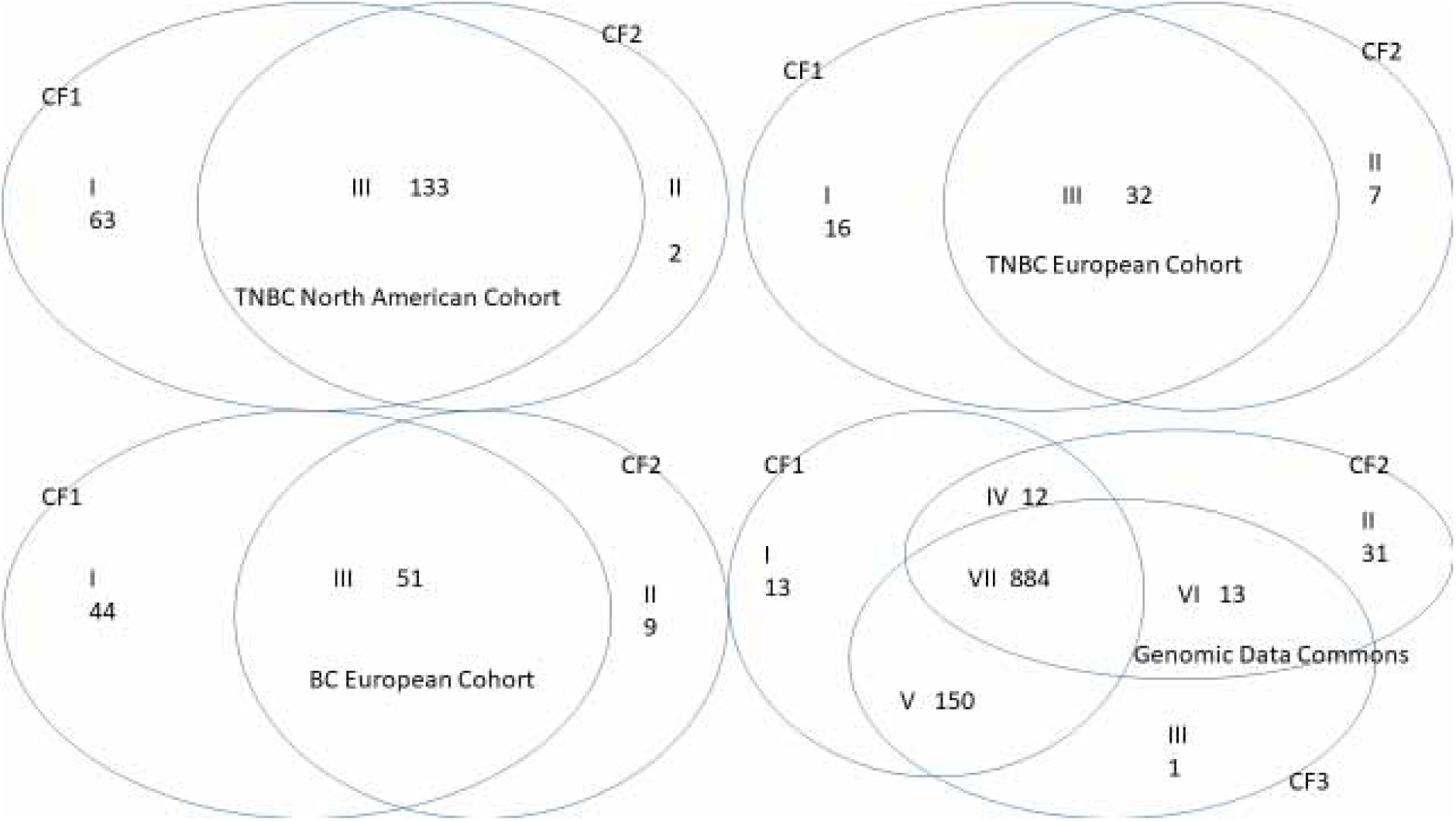
Venn diagrams of breast cancer subtypes. The first three cohorts have more than three subtypes. The fourth cohort has more than seven subtypes.

For the first TNBC (North American cohort) dataset, three genes (RNF213, RBM22, CACNG4) completely classify all 198 TNBC tumor samples into three subtypes (Figure 2) with the sensitivity of 100% and the specificity of 100%. From the individual classifiers, we can see that a decrease of RNF213 level will reduce the risk of developing TNBC, while increases in the expression levels of RBN22 and CACNG4 will reduce the risk of developing TNBC.

For the second TNBC (European cohort) dataset, four genes (MYCT1, NAT8L, NUAK2, CACNG4) completely classify all 66 TNBC tumor samples into three subtypes (Figure 2) with the sensitivity of 100% and the specificity of 100%. From the individual classifiers, we can see that a decrease in NUAK2 level will benefit the patients, while increases in the expression levels of MYCT1, NAT8L, and CACNG4 will benefit the patients. We note that there are also Her2, Luminal A and Luminal B samples in this second dataset. After adding classifier CF3: 21.8170 - 8.8170*RBM22 - 0.3047*NAT8L, all breast cancer (TNBC, Her2, Luminal A, Luminal B) patients will again be 100% accurately classified into their respective groups.

Comparing the first and second TNBC cohorts, we see that the TNBC patients from North American and the TNBC patients from European cohorts share a common gene CACNG4 and similar coefficients (−3.6692 vs. −3.5933). Otherwise, other critical genes from these two cohorts are different. This observation tells that the causes, the formations, and the therapies of TNBC can be different from region to region and race to race. We want to note that based on our knowledge in the field, there does not exist any other method that can 100% accurately classify breast cancer patients and cancer-free patients into their respective groups. With 100% accuracy, regardless of how big and how small the sample is, these genes should contain basic cancer information of TNBC disease, they should be thoroughly analyzed and explored.

On the other hand, cautions should be called with any other classifiers with lower accuracy. Using genes derived/obtained from low accuracy classifiers may lead to suboptimal results and even wrong conclusions. The formulas of these two cohorts disclose the puzzle of TNBC as gene-gene interactions and functional effects are different. Such differences can be the most important part of studying TNBC and point out new research directions for better understanding TNBC and designing better treatments.

For the third (European cohort) dataset, four genes (MYCT1, NAT8L, NUAK2, UNC5B) completely classify all 104 tumor samples into three subtypes (Figure 2) with a sensitivity of 100% and a specificity of 100%. A decrease of UNC5B level will benefit the patients in this cohort, while increases of expression levels of MYCT1 and NAT8L will benefit the patients. In addition, it can be seen that NUAK2 can benefit the patients and can also harm the patients depending on the patients’ breast cancer subtypes in Figure 2. These gene-gene relationships and genes-subtypes relationships tell efficient therapies to breast cancer patients depending on their subtypes’ determinations.

For the fourth (Genomic Data Commons) dataset, the same four genes (MYCT1, NAT8L, NUAK2, UNC5B) as for the third (European cohort) dataset completely classify 1104 tumor samples into seven subtypes (Figure 2) with the sensitivity of 100% and the specificity of 96.5%. There are 4 samples among 103 normal samples being classified as tumor samples. Note that this dataset does not offer multiple ID-ref subtypes. If genes’ expression values are taken the same as those ID-ref subtypes, the specificity may be improved to 100%. In this cohort, increases of MYCT1 and NAT8L levels can benefit the patients, while decreases of UNC5B and NUAK2 levels will benefit the patients.

Comparing the third and fourth breast cancer cohorts, the individual classifiers Data-3-CF1, Data-4-CF1, and Data-4-CF3 have the same component genes and coefficient signs. Data-3-CF2 has one more gene, NAT8L, then Data-4-CF2. However, the signs of NUAK2 coefficients in these two individual classifiers are different. We further note that to have two similar individual classifiers Data-4-CF1 and Data-4-CF3 in the final classifier is completely new in machine learning literature. These observations further reveal that breast cancer formations are more complicated than simply looking at some high/low expression values of individual genes as in the literature. The most important relations in finding critical genes linked to breast cancers are gene-gene interactions, genes-individual classifiers interactions, and their functional effects.

Comparing the second TNBC cohort, the third breast cancer cohort, and the fourth cohort, we see that increasing the levels of MYCT1 and NAT8L can benefit all patients.

In Figure 2, Venn diagrams for four different cohorts are different, with two TNBC cohorts having similar patterns, while BC European cohort and Genomic Data Commons are different. It is because the numbers of component classifiers in the final classifiers for different cohorts are different. Such phenomena tell that there are commonalities among breast cancer patients and specificities from patient to patient, i.e., the critical cancer informatics are expressed. Note that in Venn diagrams, the more intersections the groups, the more complex the disease, and the more difficult the treatment. Taking Genomic Data Commons as an example, patients in Group VII will be the most difficult to treat.

Figure 3 presents the gene-gene interactions, gene-subtype interactions, and functional effects of our identified competing classifiers. We can see clear signature patterns in each plot. This visualization tool provides a new way for breast cancer diagnosis.

**Figure 3.**
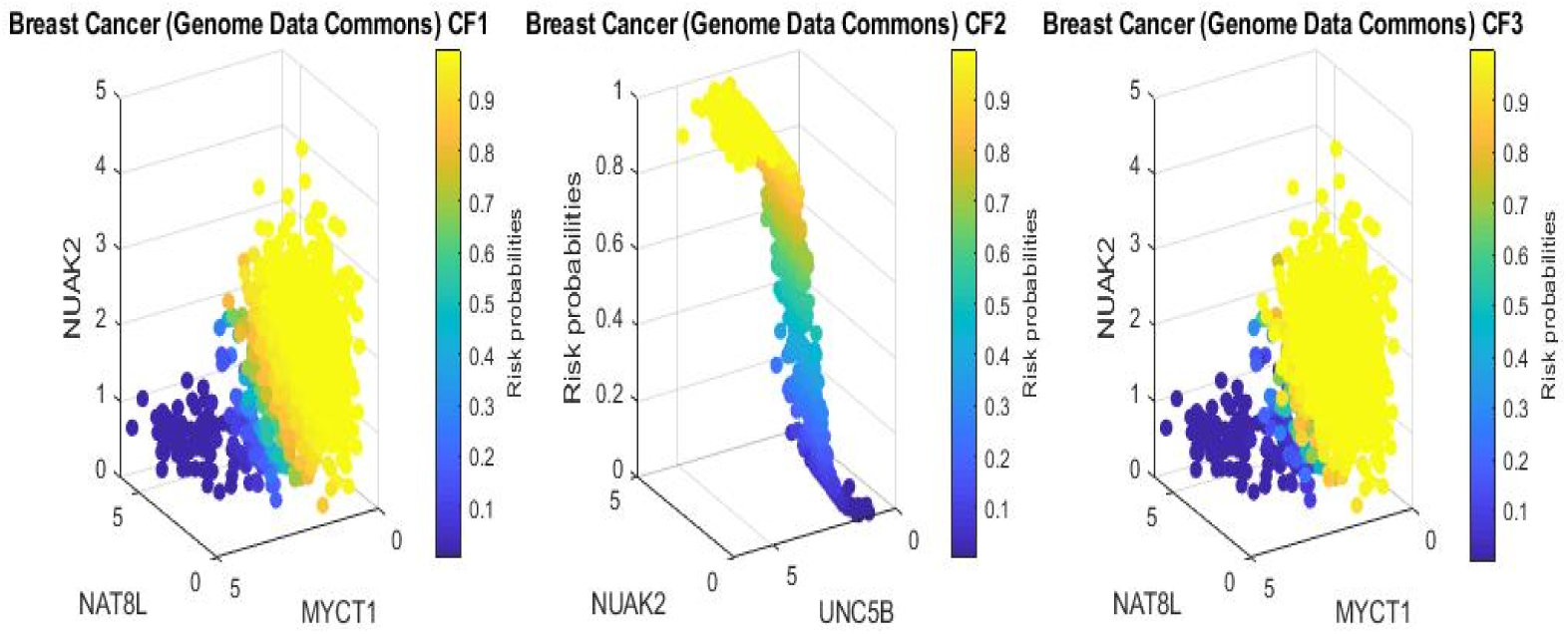
Four-dimensional plots for visualizing risk signature patterns from three competing component classifiers and the combined functional effects of gene-gene interactions and gene-subtype interactions of four genes.

### Characteristics of studying samples

All four datasets are accompanied by some characteristics of patients. Here we report their inter-relationship with the competing classifiers. Table 2 displays Sex, Age, BMI, and Grade from the first dataset (TNBC, a North American cohort). Table 3 displays Age and BMI from the second dataset (TNBC samples only, A European cohort). Table 4 is for the third dataset (breast cancer samples, A European cohort). Finally, Table 5 includes disease Stage besides Age and Sex for the fourth breast sample data set (A Genomic Data Commons – TCGA).

**Table 2.** Characteristics of the first dataset samples (TNBC, A North American Cohort)

|  | Sex |  | Age |  |  |  | BMI |  |  | Grade |  |  |
| --- | --- | --- | --- | --- | --- | --- | --- | --- | --- | --- | --- | --- |
|  | Male | Female | <=50 | (50,60] | (60,70] | >70 | Normal | Overweight | Obese | Poorly | Moderately | Well |
| CF-1 | 0 | 63 | 34 | 12 | 10 | 5 | 13 | 19 | 61 | 33 | 25 | 1 |
| CF-2 | 0 | 2 | 1 | 0 | 1 | 0 | 0 | 1 | 2 | 1 | 0 | 0 |
| CF-(1,2) | 0 | 133 | 51 | 33 | 28 | 18 | 36 | 41 | 129 | 74 | 28 | 3 |

**Table 3.** Characteristics of the second dataset samples (TNBC, A European Cohort)

|  | Age |  |  |  | BMI |  |  |
| --- | --- | --- | --- | --- | --- | --- | --- |
|  | <=50 | (50,60] | (60,70] | >70 | Normal | Overweight | Obese |
| CF-1 | 3 | 4 | 2 | 2 | 5 | 3 | 3 |
| CF-2 | 2 | 2 | 0 | 1 | 2 | 2 | 1 |
| CF-(1,2) | 9 | 10 | 5 | 1 | 12 | 6 | 6 |

**Table 4.**
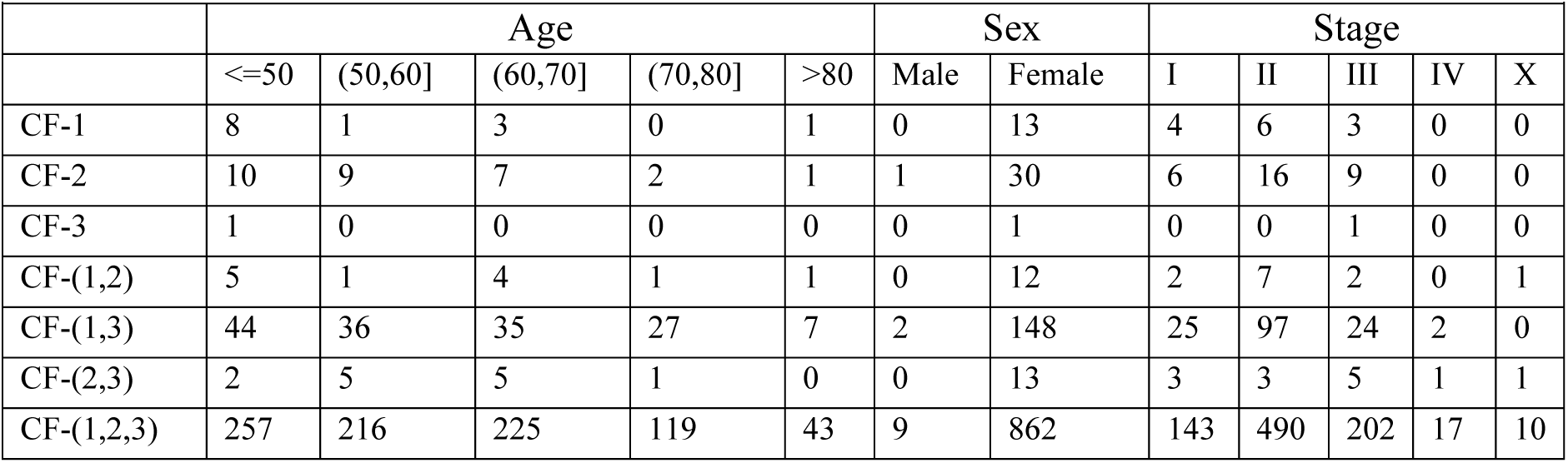
Characteristics of the third dataset samples (BC, A European Cohort)

**Table 5.**
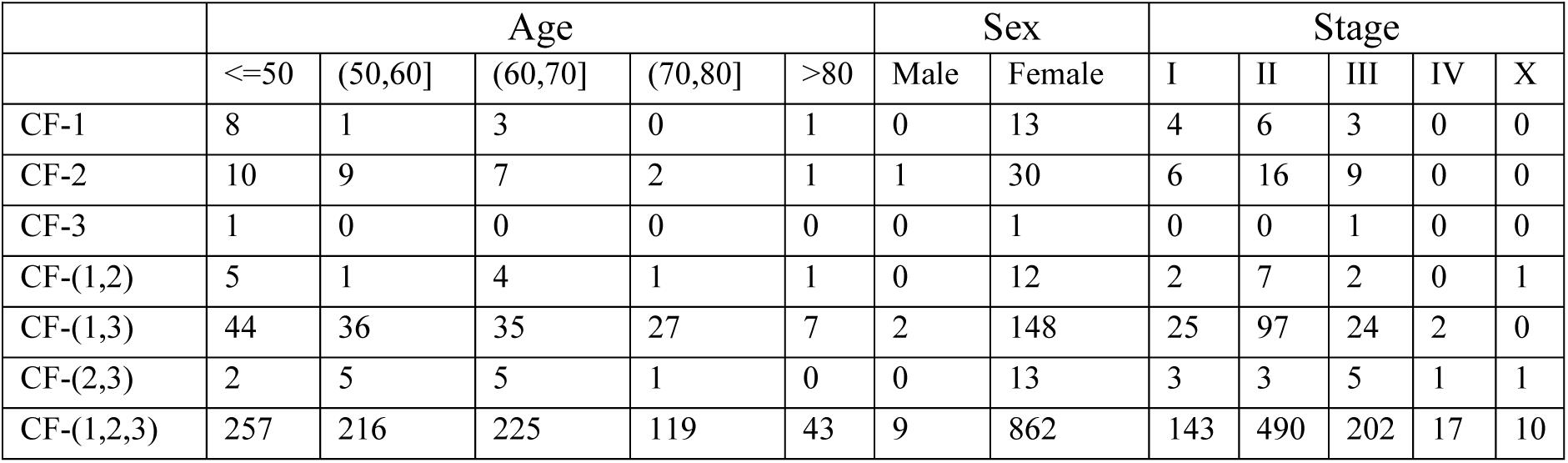
Characteristics of the fourth dataset samples (TCGA, Genomic Data Commons)

Overall, these four tables show that more patients fall in groups related to more than one competing classifier. Obese patients can make TNBC more complex. The more the competing classifiers, the worse the grade. In Table 5, Stages (IV, X) are mainly related to CF-(1,2,3), which shows the classifiers are positively correlated.

## Discussions

This study is the first time in the medical literature that breast cancer diseases can be classified almost 100% correctly using only a few (three or four) genes. The results clearly disclose the puzzle of breast cancers, including TNBC, due to the selected genes and their predicting powers through gene-gene interaction, gene-subtype interaction, and functional effects. The results also point to new treatment directions.

We note that this study does not use the primary endpoint information. It is a pure classification study. The main purpose is to identify the essential breast cancer informatics. The study has achieved 100% accuracy, which is the first in the literature. Given patients have different endpoints, the new classifier still reaches 100% accuracy, which means the classifier is robust to patients’ disease states, i.e., we can conclude that the classifier is robust regardless of primary endpoints and other individual attributes.

The discovery of critical genes can motivate many new research directions and laboratory experiments. These critical genes and their derived signature patterns (individual classifiers) can be a starting point as new biomarkers for conducting gene network analysis, testing other reported genes, and finding the causal directions of gene expression in various projects. As a result, many other existing pieces of research can be enriched. It can also be hoped that new types of diseases can be discovered. Eventually, new testing procedures and therapies for breast cancer can be designed.

These critical genes enrich the biological literature of their new functions related to breast cancer from indirect relationship to direct relationship. In many scenarios, indirect effects are more significant than direct effects as direct effects can be seen and controlled, while indirect effects are hard to see and even not to say how to control.

The risk probability of a patient developing a specific type of breast cancer in her/his life is low. Among all discovered breast cancer types, growing more than one type of breast cancer is rare. These breast cancer types compete, and one type will first be diagnosed. As a result, the competing risk factor models can efficiently model multiple breast cancer types.

This study’s inference/analysis approach can shed new light on all gene-related research, i.e., not just the breast cancer study. Researchers can apply max-linear type models in their studies. Ultimately, our new findings may make researchers’ cancer research efforts more effective and meaningful, reduce substantial research costs, and save lives and protect people.

Finally, we address an important medical practice issue. In this paper, all classifier formulas are explicitly expressed. Thus, the results in Table 1 are reproducible. Furthermore, Figures 1 and 3 show the risks of all patients. Using this paper’s results, medical doctors have a powerful tool (testing kit) in their daily work, i.e., in the diagnostic stage, diagnosing and analyzing patients’ breast cancer risks based on the four or fewer critical genes’ expression values and the computed risks; in the treatment stage, those signature patterns can be used to study the effectiveness of drugs and treatments, i.e., conduct clinical trials, e.g., survival analysis, based on classified groups; in the drug development stage, pharmaceutical companies can use the findings of critical genes to study new drugs; finally, it can be hoped that mRNA-based therapies can be introduced using the critical genes’ information in the therapy stage.

